# Interstitial leukocyte navigation through a search and run response to gradients

**DOI:** 10.1101/2021.03.03.433706

**Authors:** Antonios Georgantzoglou, Hugo Poplimont, Tim Lämmermann, Milka Sarris

## Abstract

Migrating cells must interpret chemical gradients to guide themselves within tissues. A general premise has been that gradients influence the direction of leading-edge protrusions. However, recent evidence indicates that actin flows correlate better with directed motion than protrusions. A unified chemotaxis model that integrates the role of protrusions and actin flows and that accounts for *in vivo* cell motion patterns is lacking. Here, we provide direct experimental evidence of how neutrophils sense gradients in real-time *in vivo* and demonstrate a two-stage process: first a ‘search’ phase, that requires actin network expansion by Arp2/3, whereby cells slow down, explore the environment and execute path corrections. This is followed by a ‘run’ phase, that requires Myosin-II-driven contractility and fast actin flows, whereby cells accelerate and persist in the source direction. Thus, protrusive forces support chemotaxis by enhancing signal detection, while actin flows act as the ultimate output of gradient processing.

## Introduction

Cell movement is a driving force in embryonic development, cancer cell dissemination and immune responses. To move appropriately to functional destinations, cells must accurately interpret chemical gradients in their environment. A general mechanistic principle is that gradients modulate actin polymerisation at the cell front, such that leading edge protrusions are selectively generated or stabilised in the direction of the source (‘local excitation’) (Insall, 2010; Sarris and Sixt, 2015; Van Haastert and Devreotes, 2004; Xiong et al., 2010). Concurrently, protrusions are inhibited in other parts of the cell (‘global inhibition’) through long-range diffusion of chemical factors (Franca-Koh and Devreotes, 2004) or global mechanical tension (Diz-Muñoz et al., 2016; Houk et al., 2012), establishing or reinforcing cell polarity towards the gradient direction. In this framework, protrusive forces are the main output of chemoattractant gradient interpretation.

While the influence of chemical gradients on protrusions has been observed in many cells, it remains unclear whether this mechanism alone is sufficient to account for directed motion and the complex motion patterns of cells *in vivo*. Leading edge expansion is driven by branching of actin networks through the actin nucleator Arp2/3 (Rotty et al., 2013). Yet, accumulating evidence indicates that loss of Arp2/3 does not compromise chemotaxis in a direct fashion. In cells moving in a mesenchymal-like fashion, like fibroblasts, Arp2/3 was found to be dispensable for persistent protrusions and directed motion, unless the chemical gradient was immobilised on a substrate (haptotaxis) (Wu et al., 2012). As mesenchymal locomotion relies on adhesion of protrusions to the extracellular matrix (Case and Waterman, 2015), this suggested that leading edge expansion is important for coordinating optimal adhesions with the substrate rather than locomotion *per se*. In cells moving in an amoeboid fashion, such as leukocytes, organised focal adhesions are dispensable and, under confinement, cells can move in the absence of dedicated adhesion molecules (Lämmermann and Sixt, 2009; Paluch et al., 2016). Instead, they use flexible deformations and friction against the confining substrate to swiftly squeeze through tissue interstices (Lämmermann and Sixt, 2009; Paluch et al., 2016). Arp2/3 was found to be dispensable for neutrophil and dendritic cell motility in 1D and 2D substrates (Leithner et al., 2016; Vargas et al., 2016; Wilson et al., 2013) but more specifically required for fast movement through dense 3D matrices (Leithner et al., 2016). These observations suggest that protrusion expansion in amoeboid cells is important for exploration and discovery of optimal paths of least resistance rather as a driving force of movement. If protrusion extension is not a primary driver of cell locomotion, then how do chemical gradients redirect motion?

Actin flows have been proposed as a universal driving force for cell locomotion. While actin polymerisation at the cell front is translated to protrusions to a certain extent, cell contractility produces membrane tension that counteracts the anterograde force of growing actin networks, resulting in retrograde flow of actin filaments (Callan-Jones and Voituriez, 2016; Case and Waterman, 2015). This generates a retrograde force that, when transduced to the substrate, can translocate the cell forward (Callan-Jones and Voituriez, 2016; Case and Waterman, 2015). Force transduction can be achieved by specialised adhesion molecules, notably integrins, when cells move along 2D surfaces, or through friction, when cells are moving in confined spaces (Paluch et al., 2016). Accordingly, the speed of retrograde actin flow is a direct determinant of cell speed (Maiuri et al., 2015). Interestingly, retrograde actin flows also appear to play a role in cell directionality and chemoattractant signalling. First, actin flows reinforce polarity and persistence of motion, a relationship that is thought to arise from transport of polarity factors by retrograde flows (Lange et al., 2016; Maiuri et al., 2015). Secondly, the coordination of actin flows in different parts of the cell correlates with directional and persistent motion (Yolland et al., 2019). Finally, the speed of actin flows can be directly influenced by the concentration of chemoattractant in the environment (Hons et al., 2018). Together, these observations suggest that actin flows could play a key part in gradient interpretation.

Currently, we are lacking a unified and physiologically relevant mechanistic model for chemotaxis. Models thus far do not integrate the role of protrusions and actin flows and are essentially based on *in vitro* observations. How gradients affect sub-cellular scale cytoskeletal dynamics to generate complex cell motion patterns at the tissue scale is still unexplored experimentally. Here, we address this problem by exploiting zebrafish and mouse neutrophils as a model, acute *in vivo* chemotaxis assays and quantitative imaging. We show that cells respond to a newly encountered gradient in two key stages: First a ‘search’ phase, during which wandering neutrophils abruptly decelerate and actin polymerisation is stimulated at the cell front in an Arp2/3-dependent manner. This phase is required for signal exploration and correct turning in the direction of the source. Next, cells switch into ‘run’ phase, whereby Myosin-II contractility promotes fast actin flows, high speed and directional persistence. Thus, leading edge expansion and actin flows are both involved in chemotaxis but drive distinct parts of the gradient interpretation process, with protrusive forces driving the initial search and contractile forces driving the ultimate output of persistent motion.

## Results

### Zebrafish and mammalian neutrophils respond to gradients via small turns and an increase in directional speed and persistence

To identify conserved effects of gradients on leukocyte motion patterns *in vivo*, we exploited a two-photon laser wound assay in zebrafish and mouse tissue. This provides acute activation of endogenous chemical gradients in a tissue and thus enables a direct comparison of cell behaviour before and after gradient exposure. We first examined neutrophil gradient responses in zebrafish. We performed laser wounds and imaging by two-photon microscopy in the ventral fin of 3 days-post fertilisation (dpf) transgenic larvae that express a fluorescent probe in neutrophils (Figure 1A, Supplementary Figure 1A and Video 1). The ventral fin is devoid of neutrophils but located adjacent to the caudal hematopoietic tissue (CHT), a site of neutrophil residence and accumulation. We used a procedure that allowed us to collect data on neutrophil motion both before and after wounding in the same tissue (Supplementary Figure 1A). Initially, we forced neutrophil exit from the CHT by adding the chemoattractant Leukotriene B4 (LTB4) in the medium (Coombs et al., 2019). This stimulates mobilisation of neutrophils from the CHT into the ventral fin, where the cells remain highly motile but responsive to secondary gradients by subsequent wounding (Coombs et al., 2019). After recording neutrophil motion for approximately 30 minutes, we performed an acute laser wound. This led to rapid redirection of cell motion towards the wound focus (Figure 1A and Video 1).

**Figure 1.**
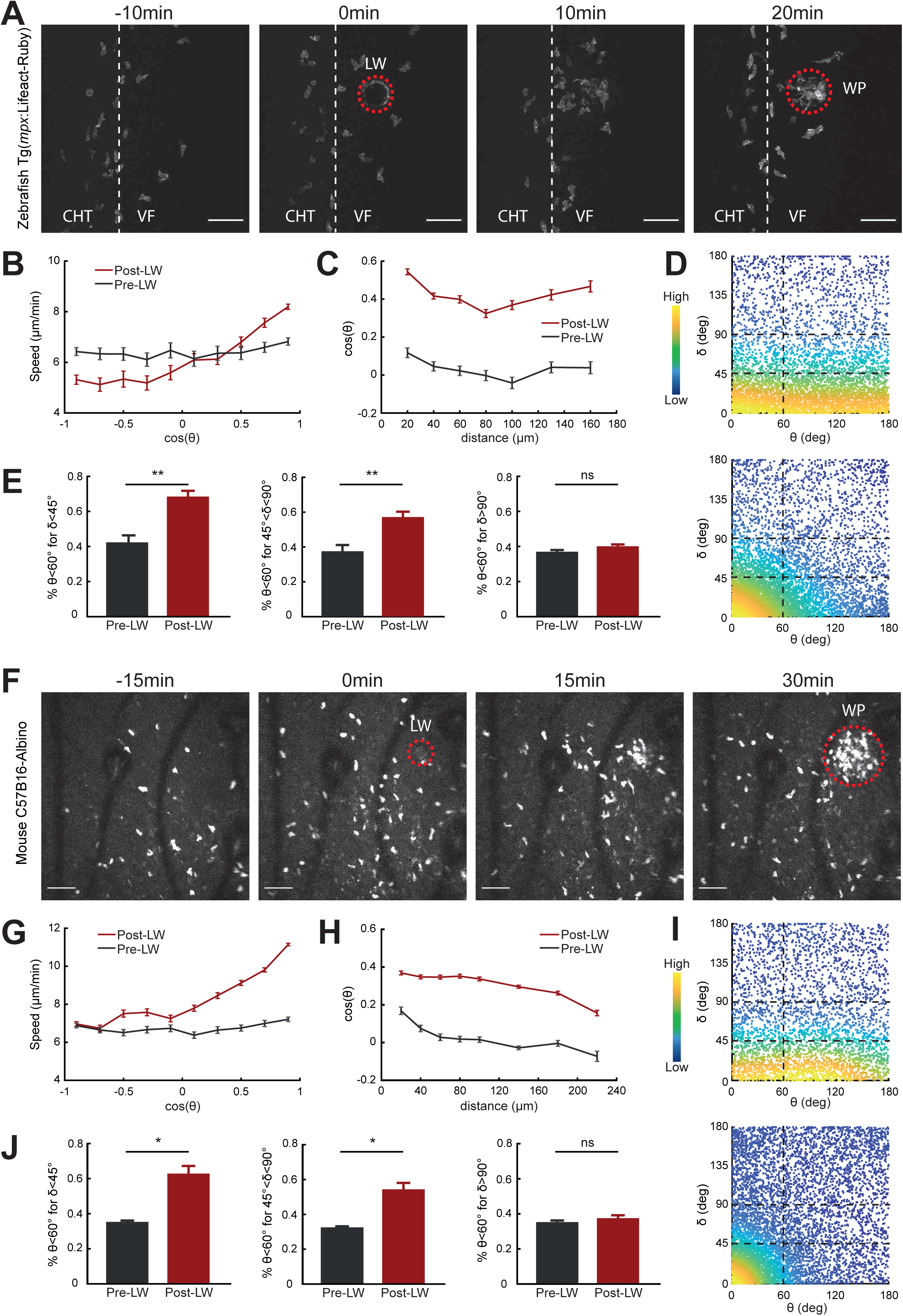
Gradients cause a directional bias on speed and persistence and small reorientations towards the source. Analysis of neutrophil migration responses to gradients in zebrafish (A-E) and mice (F-J). (A,F) Time-lapse sequence of two-photon confocal image projections, showing fluorescent neutrophils in 3 dpf Tg(*mpx:*Lifeact-Ruby) transgenic larvae (A) and CMTPX dye-labelled neutrophils (white) in a C57B16-Albino mouse (F), migrating pre- and post-laser wound (LW - red dashed line). LW occurs at 0 min. Scale bar = 50μm. (B,G) Cell speed in relation to the cosine of theta for pre- and post-laser wound. (B) n=343-1283 cell-steps per bin pre-wound, n=236-3168 cell-steps per bin post-wound. (C,H) Cell cosine of theta in relation to the distance from the closest point of the wound perimeter (WP). (C) n=517-863 cell steps per bin for pre-wound, n=444-1272 cell steps per bin for post-wound. (D,I) Scatter plot of cell delta in relation to theta, colour-coded for the density of points, for pre-laser wound (top) and post-laser wound (bottom), for zebrafish (D) and mice (I). Black dashed lines indicate the groups of data points taken for analysis for (E,J). (E,J) Percentage of cell persistent steps (delta < 45°), small turns (45° < delta < 90°) and large turns (delta > 90°), respectively, with theta < 60°, for zebrafish (E) and mice (J). Wilcoxon matched-pairs signed rank test. (B,C,E) Data from 12 larvae. (G) n=629-4879 cell steps per bin for pre-wound, n=2152-6673 cell steps per bin for post-wound. (H) n=1145-3156 cell steps per bin for pre-wound, n=1580-12000 cell steps per bin for post-wound. (I) Plotted data from 2 representative mice. (G,H,J) Data from 7 mice.

To diagnose the effects of gradients on neutrophil behaviour, we performed trajectory analyses. Previous studies indicated that cells moving in exogenous, constitutive gradients have biased speed in the direction of the source (Sarris et al., 2012). To explore whether acute endogenous gradients bias speed according to orientation, we plotted instantaneous speed against the cosine of the approach angle θ (theta), which represents the angle of the motion vector in relation to the closest point of the wound perimeter (Figure 1B and Supplementary Figure 1B). We found that neutrophils increased speed towards the source and decreased speed in the opposite direction, in contrast to their movement prior to gradient exposure, where speed fluctuated independently of motion direction. To quantify global changes in orientation, we plotted the cosine of the approach theta across different distances from the closest point of the wound perimeter. This revealed a marked increase in orientation bias across a distance range as far as 160µm from the wound perimeter (Figure 1C).

Quantification of average orientation of the cells does not distinguish whether neutrophils are performing oriented turns or changing their persistence. To distinguish these effects we introduced another metric, angle δ (delta), which represents the change in angle between successive steps in neutrophil motion (Supplementary Figure 1B). We distinguished a range of delta: persistent steps (defined here as delta < 45°), followed by small turns (defined here as delta in region 45-90°) and large U-turns (defined here as delta in region 90-180°) (Figure 1D). Prior to gradient exposure, theta were distributed in an isotropic fashion across the range of delta, indicating no orientation bias in any of these steps (Figure 1D). Post-gradient exposure, we observed a significant increase in persistent steps and small turns but no increase in large turns (Figure 1D,E). This is consistent with the idea that polarity has a certain memory and pre-polarised cells are resistant to polarity reversal (Albrecht and Petty, 1998; Arrieumerlou and Meyer, 2005; Gerisch and Keller, 1981; Skoge et al., 2014).

To test the conservation of these gradient effects in mammalian neutrophils, we analysed data of two-photon imaging and laser wounding in mouse ear skin tissue using intravital microscopy (Figure 1F and Supplementary Figure 1C). We injected CellTracker Red CMTPX dye-labelled neutrophils, that were isolated from bone marrow of C57Bl6 mice, into the ear dermis of C57Bl6-Albino mice and waited for 4 hours before intravital imaging was started. Previous studies have shown that neutrophil dynamics in this adoptive transfer are comparable to the wound dynamics of endogenous mouse neutrophils (Lämmermann et al., 2013). Once imaging started, the motion pattern of neutrophils was recorded for 30 minutes before and after laser wounding. We observed marked chemotaxis and accumulation of neutrophils at the wound (Figure 1F and Video 2). Consistent with observations in zebrafish, neutrophils increased their directional speed and the probability of persistent steps and small turns in the direction of the source (Figure 1G-J).

Thus, neutrophils respond to newly encountered gradients *in vivo* by small adjustments to their migration path and a marked change in speed and persistence in the source direction.

### Small turns are associated with front enrichment of actin polymerisation probes and slow motion

To relate gradient-induced motion patterns to cytoskeletal dynamics, we analysed actin dynamics in transgenic Tg(*mpx*:Lifeact-Ruby) zebrafish larvae (Yoo et al., 2010), whose neutrophils express Lifeact, a 17-amino-acid peptide that binds actin microfilaments (Riedl et al., 2008). Using the same two-photon laser wound assay, we followed the migration of neutrophils before and after wounding. We generated algorithms to automatically define the front and rear portions of the cells, based on the direction of motion and the geometrical centre of the cells. We then computed actin polarity as the ratio of fluorescence intensity at the front versus rear of the cell. Using cross-correlation analysis, we found that phases of slow motion strongly correlated with front enrichment of Lifeact and, conversely, phases of fast motion correlated with rear enrichment of Lifeact (Figure 2A-D,F). This correlation was observed both before and after laser wounding, suggesting that rear Lifeact configuration generally corresponds to phases of fast movement. To relate actin dynamics to cell directionality, we performed similar correlation analyses of actin polarity against the delta metric (angle in relation to previous step, whereby small angle indicates persistent movement). Interestingly, while before laser wounding there was no clear correlation between directionality and Lifeact distribution, post-wounding we detected a correlation between front Lifeact enrichment with cell turning (and thus of rear Lifeact enrichment with persistent steps) (Figure 2E,G).

**Figure 2.**
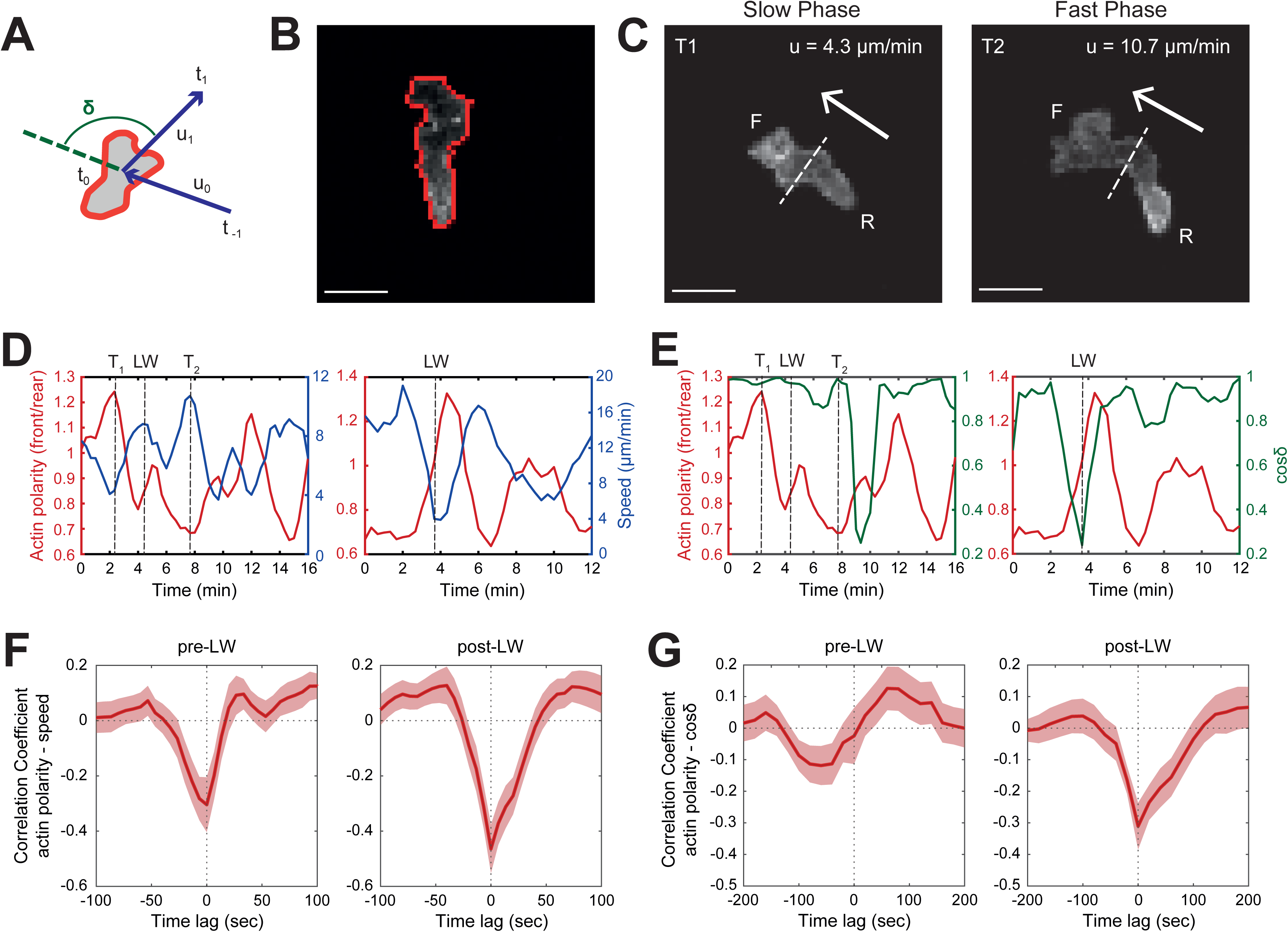
Relationship of motion patterns with distribution of actin polymerisation probes. (A) Scheme of a neutrophil, indicating the definition of angle δ (delta), which is the angle between the velocity vectors between three successive time-points. (B) Outline (red line) of an example of segmented neutrophil within a Tg(*mpx:*Lifeact-Ruby) transgenic larva. Scale bar = 10μm. (C) Example of a neutrophil in slow (T_1_) and fast (T_2_) phase of motion, with an indication of cell speed in the same time point. Arrow indicates the direction of motion. Dashed lines indicate the automated separation of the front (F) and rear (R) part of cell. Scale bar = 10μm. (D) Example of time evolution of actin polarity (left axis - red colour) and speed (right axis - blue colour) from two individual neutrophils. (E) Example of time evolution of actin polarity (left axis - red colour) and cosine delta (right axis - green colour) from two individual neutrophils. (D,E) T_1_ and T_2_ correspond to the time-points of slow and fast motion, respectively, that are shown in C. LW denotes the time-point of the laser wound. (F) Temporal cross-correlation between actin polarity and speed, pre- (left) and post-laser wound (right). (G) Temporal cross-correlation between actin polarity and cosine of delta, pre- (left) and post-laser wound (right). (F,G) Average from n=21 migrating cells, from 9 larvae. Mean and SEM are shown.

Together, this analysis revealed that front Lifeact enrichment is associated with stages when cells slow down and make directional changes.

### Fast persistent motion is associated with rear enrichment of actin polymerisation probes and fast actin flows

Front Lifeact enrichment is indicative of actin polymerisation in this area and expansion of the leading edge (Riedl et al., 2008). However, the interpretation of rear Lifeact enrichment during fast phases of motion remained less clear. Recent studies have suggested that in fast-moving cells, Lifeact can accumulate at the rear as a result of advection of the probe by fast actin flows (Yamashiro et al., 2019). Accordingly, the rear Lifeact enrichment we observed could have reflected phases of accelerated actin flows, as opposed to new actin network formation at the rear. To test the relationship of actin flows to neutrophil motion *in vivo*, we performed spinning disk confocal microscopy imaging with transgenic Tg(*lyz*:DsRed2)^nz50^/Tg(*actb1*:myl12.1-eGFP) (Behrndt et al., 2012; Hall et al., 2007) zebrafish larvae, expressing a broad expression of non-muscle Myosin-II (*myl12.*1) as well as a neutrophil DsRed marker. Myosin-II-GFP binds actin microfilaments with high affinity, permitting the tracking of actin flows (Maiuri et al., 2015). We utilised high-resolution imaging through spinning disk confocal microscopy coupled to a mechanical wound assay (tail-fin amputation) to follow the migration of neutrophils towards the tail fin within 1-2 hours post-wounding using high temporal resolution (Supplementary Figure 1D -left panel, Figure 3A and Video 3).

**Figure 3.**
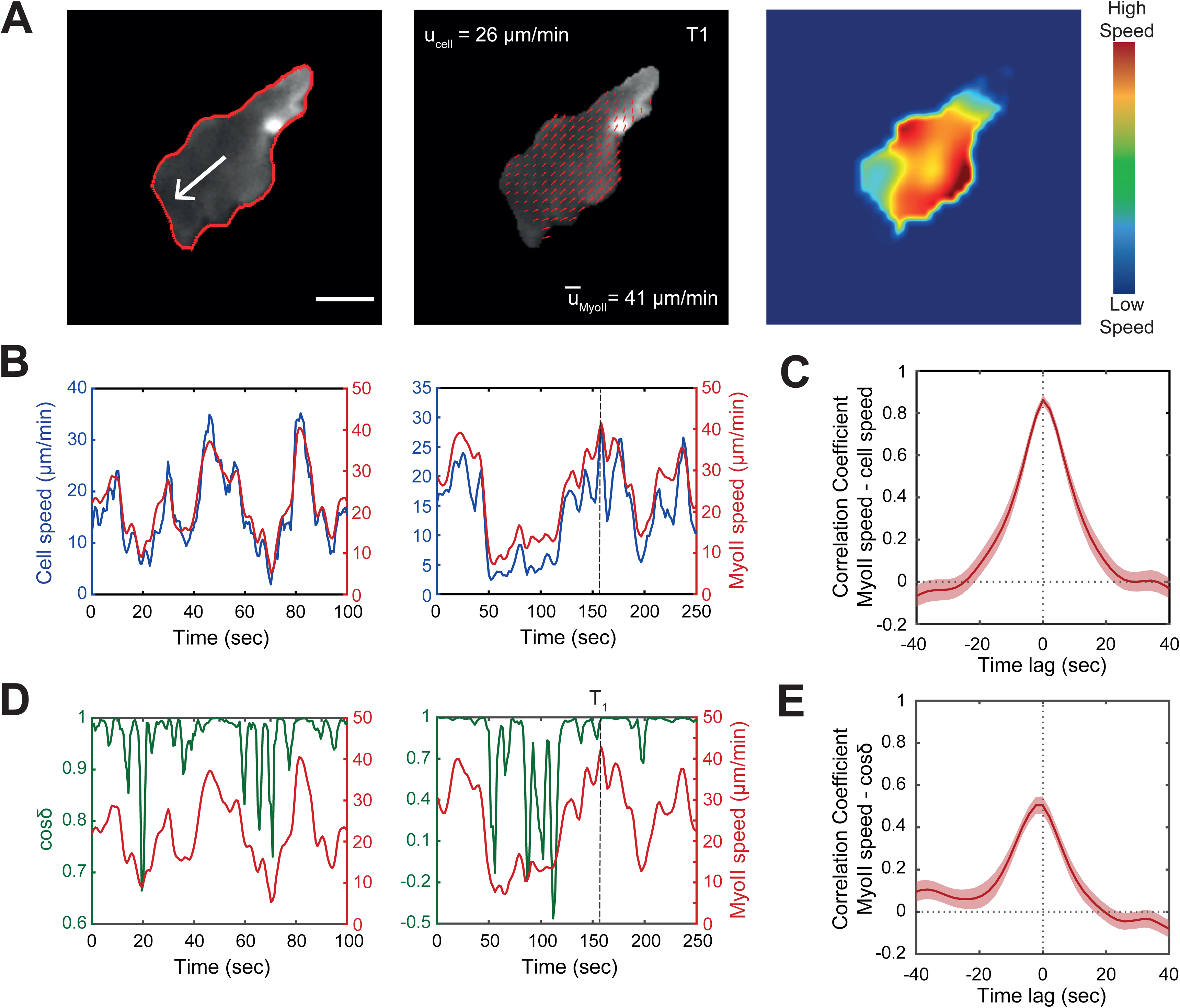
Relationship of motion patterns with speed of actin flows. (A,B,C) Left: outline (red line) of a representative segmented neutrophil within a Tg(*actb1*:myl12.1-eGFP) transgenic larva. White arrow indicates the vector of speed. Scale bar = 5μm. Middle: instantaneous Myosin-II (MyoII) retrograde flow. Red arrows indicate the velocity vector fields of Myosin-II retrograde flow. U_cell_ is the instantaneous speed of the cell. U_MyoII_ is the mean speed of the Myosin-II retrograde flow. Right: heatmap of the speed of the Myosin-II retrograde flow. Colour-bar indicates low- and high-speed values, respectively. (B) Examples of time evolution of neutrophil speed (blue) and Myosin-II retrograde flow speed (red) from two individual neutrophils. T_1_ shows the time-point that corresponds to time-frame (A). (C) Temporal cross-correlation between Myosin-II retrograde flow speed and neutrophil speed. (D) Examples of time evolution of cosine delta (green) and Myosin-II retrograde flow speed (red) from two individual neutrophils. T_1_ shows the time-point that corresponds to time-frame (A). (E) Temporal cross-correlation between Myosin-II retrograde flow speed and neutrophil cosine of delta. (C,E) n=25 cells, from 6 larvae. Mean and SEM are shown.

To track retrograde actin flows, we used Particle Image Velocimetry (PIV) (Figure 3A and Video 3) (Betz et al., 2009; Davis et al., 2015). To quantify speed of retrograde flow in the reference system of the moving cell, we subtracted the centroid of surface-segmented neutrophil of every time-frame from the centroid of the first time-frame (Video 3). Plotting the speed of actomyosin flow against the speed of the cell across time for individual cells, suggested a close correlation between the two parameters (Figure 3B). Cross-correlation analysis across many cells revealed that the speed of actomyosin flows in relation to the cell was tightly correlated to the speed of motion (Figure 3C). Conversely, the speed of actin flows was inversely correlated with cell turning (large delta) (Figure 3D,E). Thus, fast phases of motion and rear Lifeact configuration are associated with persistent directionality and fast actin flows that is consistent with the observed rear distribution of actin probes.

### Neutrophils respond to gradients through a stepwise sequence of actin dynamics

Thus far we had identified global gradient-induced motion patterns at the tissue-scale and associated subcellular actin dynamics. It remained unclear how neutrophils respond to gradients precisely at the moment of gradient exposure and how relevant cytoskeletal dynamics evolve in time. To establish this, we profiled time-resolved changes in actin dynamics immediately before and after gradient exposure (Figure 4, Video 4). This revealed that when neutrophils first experience the gradient, within the first 2-4 minutes, they decelerated, concomitantly with a peak in front actin enrichment (Figure 4A-B, Video 4). Towards the end of this phase, neutrophils completed a small turn in the direction of the source (change in delta and theta) (Figure 4B). Subsequently, cells re-initiated movement in a non-synchronised manner; such asynchrony may relate to the time needed to encounter signal and negotiate the direction of leading-edge protrusions. For the analysis, we synchronized cells to the timepoint of initiation of movement and we found that Lifeact was re-localised towards the rear of the cell concomitant with acceleration of cell motion and maintained persistent direction (stable delta) (Figure 4C, Video 4). When quantifying the persistence of motion during a time-window of 3 and 6 minutes before versus after the beginning of movement, we found that cells increased their persistence of movement after gradient exposure (Figure 4D). Thus, we can distinguish two stages in the neutrophil gradient response: a first stage with active leading edge actin dynamics and deceleration of motion, during which directional changes take place, and a second phase of fast actin flows and persistent and rapid motion.

**Figure 4.**
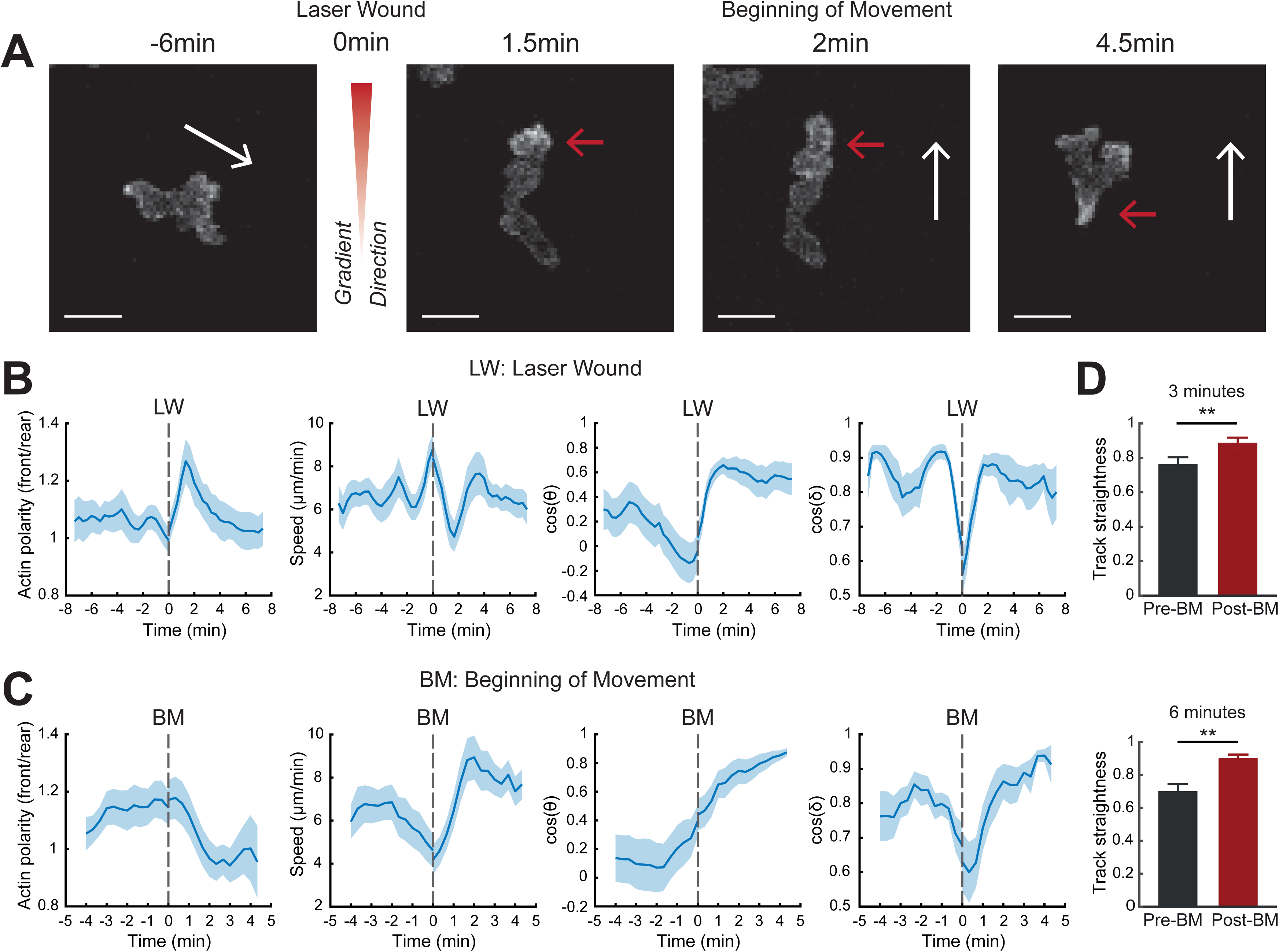
A two-stage response to newly encountered gradients. (A) Time-lapse sequence of two-photon confocal image projections showing a neutrophil (white) in a Tg(*mpx*:Lifeact-Ruby) zebrafish larva, at 6 min pre-laser wound and at 1.5 min, 2 min and 4.5 min post-laser wound. Cell movement started at 2 min post-laser wound for this example cell. White arrows indicate the speed vector. Red arrows indicate the side of cell with higher Lifeact intensity. Scale bar = 10μm. (B, C) Neutrophil actin polarity, speed, cosine of theta and cosine of delta, in relation to time, respectively. Time-sequence was synchronised based on time of the laser wound (B) or the time that each individual neutrophil beginning of movement post-laser wound (C). (B) n=21 cells, from 9 larvae. (C) n=18 cells, from 9 larvae. (D) Neutrophil track straightness, for 3 minutes (top) and 6 minutes (bottom) respectively, pre- and post-cell beginning of movement (BM). n=18 cells, from 9 larvae. Mean and SEM are shown. Wilcoxon matched-pairs signed rank test.

### Directional turns during the first phase of chemotaxis involve a searching signalling pattern

The first phase of gradient sensing was characterised by front actin enrichment and culminated in small turns in direction of movement. According to *in vitro* data, this can either result from purely spatial sensing of the gradient, i.e. resolution of the gradient independently from actin dynamics and motion, or from exploratory ‘pseudo-spatial’ sensing that requires feedback from cell exploration (Insall, 2010; Parent et al., 1998; Parent and Devreotes, 1999; Sarris and Sixt, 2015; Servant et al., 2000). A key measure for this is to test whether signalling mediators downstream of chemoattractant receptors, such as the PI3K product PIP3, can polarise in the direction of the source in the absence of active exploration (Parent et al., 1998; Servant et al., 2000). To establish this for the first time in a live tissue context, we assessed the response of the signalling mediator PI3K after inhibition of actin polymerisation with Latrunculin-B (LatB) (Parent et al., 1998; Servant et al., 2000). We introduced LatB in the medium of transgenic Tg(*mpx*:PHAKT-EGFP) (Yoo et al., 2010) zebrafish larvae (whereby PH-AKT is a probe for the PI3K product PIP3) after mechanical wounding and after the migration of neutrophils in this area had begun (Supplementary Figure 1D and Video 5A). Within 20 minutes, neutrophils were immobilised and many of them lost their polarised shape and became more rounded (Supplementary Figure 2A and Video 5A). Subsequently, we added LTB4 in the zebrafish medium to generate new outward chemical gradients in the ventral fin to assess the signalling response of the immobilised cells (Supplementary Figure 1D). We analysed the distribution pattern of PIP3 before and after gradient exposure. We found that the standard deviation and contrast of PIP3 fluorescence intensity increased in the first few minutes after gradient exposure suggesting that the gradient induced fluctuations of PIP3 intensity in the cells (Supplementary Figure 2A-C). However, the mean intensity was not increased and the orientation of PIP3, quantified as the ratio of fluorescence in the part of the cell facing the source versus the opposite direction, was on average not biased towards the source direction (Supplementary Figure 2A-C).

To complement these observations with endogenous gradients, we used LatB inhibition in neutrophils constitutively migrating in an unspecific fashion in the head (Supplementary Figure 2D and Video 5B). Upon immobilisation of the cells, we performed a laser wound, which leads to quick release of primary attractants, such as fMLP or ATP, immediately after wounding (Futosi et al., 2013; Kolaczkowska and Kubes, 2013; Poplimont et al., 2020). Analysis of PIP3 fluctuations before and after wounding again indicated that the standard deviation and contrast of PIP3 fluorescence intensity significantly increased, but not the mean intensity (Supplementary Figure 2E-F).

Together, these results suggested that, in the absence of actin dynamics, gradients induce a searching signalling pattern and that directional decisions during the first phase of chemotaxis rely on active exploration.

### Differential roles of protrusive and contractile forces in ‘search and run’ response to gradients

As our analysis divides gradient sensing into two stages, a searching phase and a running phase, we then investigated the role of protrusive and contractile forces in each phase. To address this, we visualised neutrophil motion and actin dynamics in transgenic Tg(*mpx*:Lifeact-Ruby) zebrafish larvae before and after gradient exposure in the presence of the Arp2/3 inhibitor CK666 and the Myosin-II inhibitor blebbistatin, which inhibit protrusive and contractile forces respectively (Figure 5A-C and Video 6). We first exposed larvae to LTB4, to stimulate mobilisation of neutrophils in the ventral fin and unspecific motility in this tissue followed by laser wounding to capture redirection of motion towards a new acute gradient. Co-application of CK666 led to a reduction of cell speed (Supplementary Figure 3A,B), consistent with previous observations in dendritic cells within dense matrices (Leithner et al., 2016). However, a directional bias on cell speed was still preserved in CK666-treated cells (Supplementary Figure 3B). Blebbistatin-treated cells showed reduced cell speed but also a loss in directional bias on speed (Supplementary Figure 3A,C). These cells were characterised by an overdeveloped leading edge and defects in uropod retraction (Figure 5C and Video 6C), as previously reported (Yoo et al., 2010). In the first phase of chemotaxis, while control DMSO-treated and blebbistatin-treated cells showed front actin enrichment and oriented turns, these events were suppressed in CK666-treated cells (Figure 5D-G). This was accompanied by suppressed turning during this phase, as the cosine of theta showed limited increase after gradient exposure in comparison to DMSO-or blebbistatin-treated exposed cells (Figure 5D-F). Consistent with this, we found a reduced probability of small turns in population-based trajectory analyses in CK666-treated cells in comparison to DMSO or blebbistatin-treated cells (Figure 5G). Consistent with shared defects in cell speed (Supplementary Figure 3), both blebbistatin and CK666-treated cells did not show rear Lifeact enrichment at the beginning of cell movement, suggesting reduced speed of actin flows in these cells (Figure 5H-J). On the other hand, blebbistatin-treated cells showed particular defects in persistence of orientation, as indicated by the fluctuations in cosine of delta within the first 6 minutes post beginning of movement (second phase of chemotaxis) (Figure 5H-J). Specifically, DMSO and CK666-treated cells maintained persistent motion in the first 3 minutes after beginning of neutrophil movement, whereas the orientation of blebbistatin-treated cells was more variable (Figure 5H-J). Quantification of cell motion persistence before and after the beginning of movement indicated a specific defect in blebbistatin-treated cells at a time scale of 3 minutes (Figure 5K). CK666-treated cells also showed reduced persistence at a time scale of 6 minutes, but to a lesser extent than blebbistatin-treated cells (Figure 5K).

**Figure 5.**
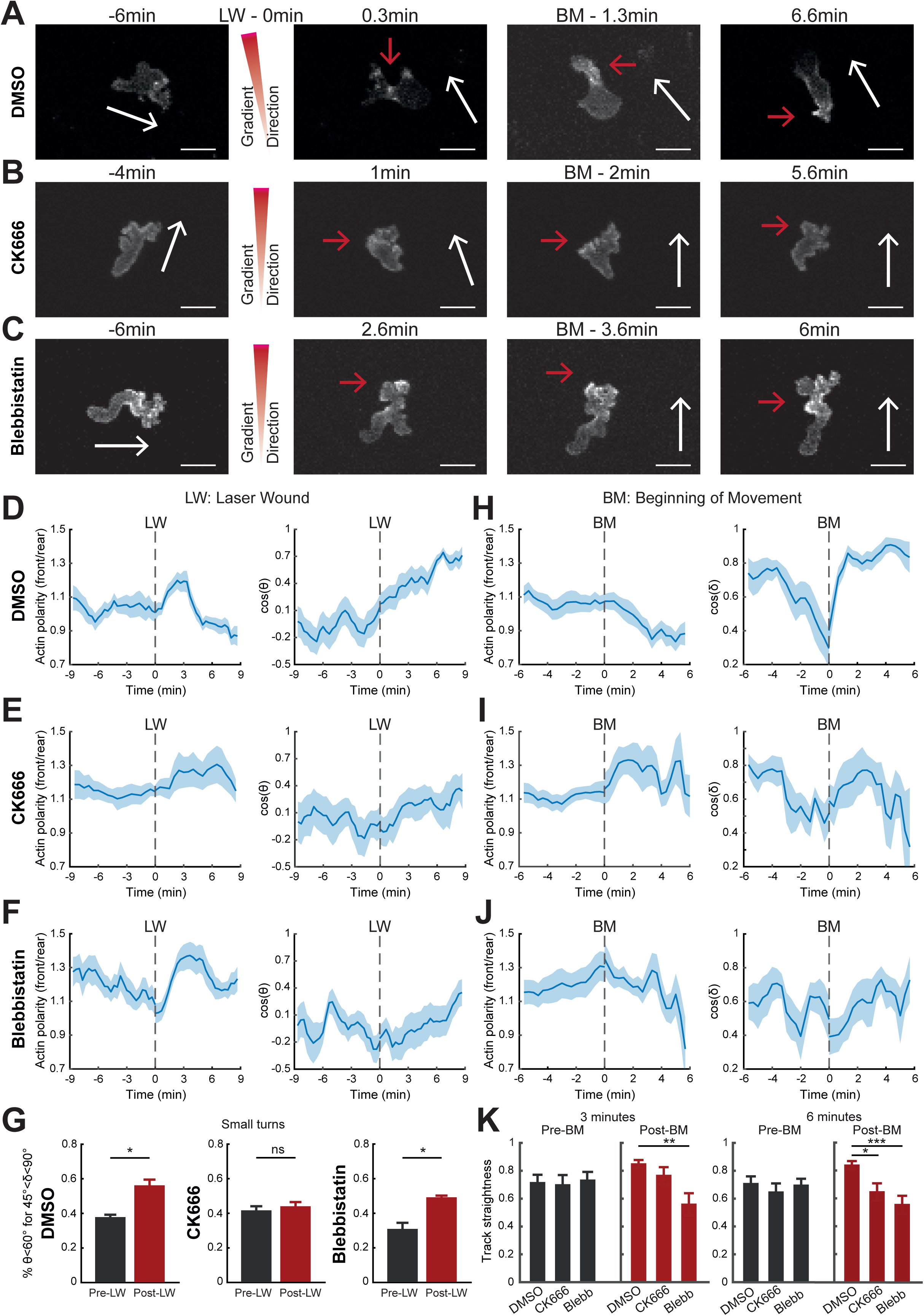
Differential contributions of Arp2/3 and Myosin-II in gradient responses. (A,B,C) Two-photon time lapse image projections showing neutrophils in Tg(*mpx:*Lifeact-Ruby) zebrafish larvae treated with control DMSO (A), CK666 (B) and blebbistatin (C). White arrows indicate the speed vector. Red arrows indicate the side of neutrophil with higher Lifeact intensity. BM: beginning of movement. Scale bar = 10μm. (D,E,F) Neutrophil actin polarity (left) and cosine of theta (right) in relation to time for DMSO (n=17 cells from 8 larvae) (D), CK666 (n=11 cells from 5 larvae) (E) and blebbistatin (n=13 cells from 4 larvae) (F). Time-sequence was synchronised based on time of the laser wound (LW - 0 min). (G) Percentage of cells with persistent steps (theta < 60° for delta < 45°). n=9 larvae for DMSO, n=5 larvae for CK666 and n=4 larvae for blebbistatin. Wilcoxon matched-pairs signed rank test. (H,I,J) Neutrophil actin polarity (left) and cosine of delta (right) in relation to time for DMSO (H), CK666 (I) and blebbistatin (J). Time-sequence was synchronised based on time of the neutrophil beginning movement (BM - 0 min). (K) Track straightness for 3 minutes (left) and 6 minutes (right) post-neutrophil movement, for DMSO, CK666 and blebbistatin. One-way ANOVA with Dunnett’s multiple comparisons test. (H-J) n numbers same as for D-F. Mean and SEM are shown.

Together these analyses indicated that the ‘search and run’ response to chemical gradients requires coordinated contribution of protrusive and contractile forces. Protrusive, exploratory structures govern the first phase of the chemotaxis response and enable renegotiations of the migration path whereas contractility is important in the second phase of the gradient response to promote fast actin flows that in turn accelerate and reinforce motion in the right direction (Figure 6).

**Figure 6.**
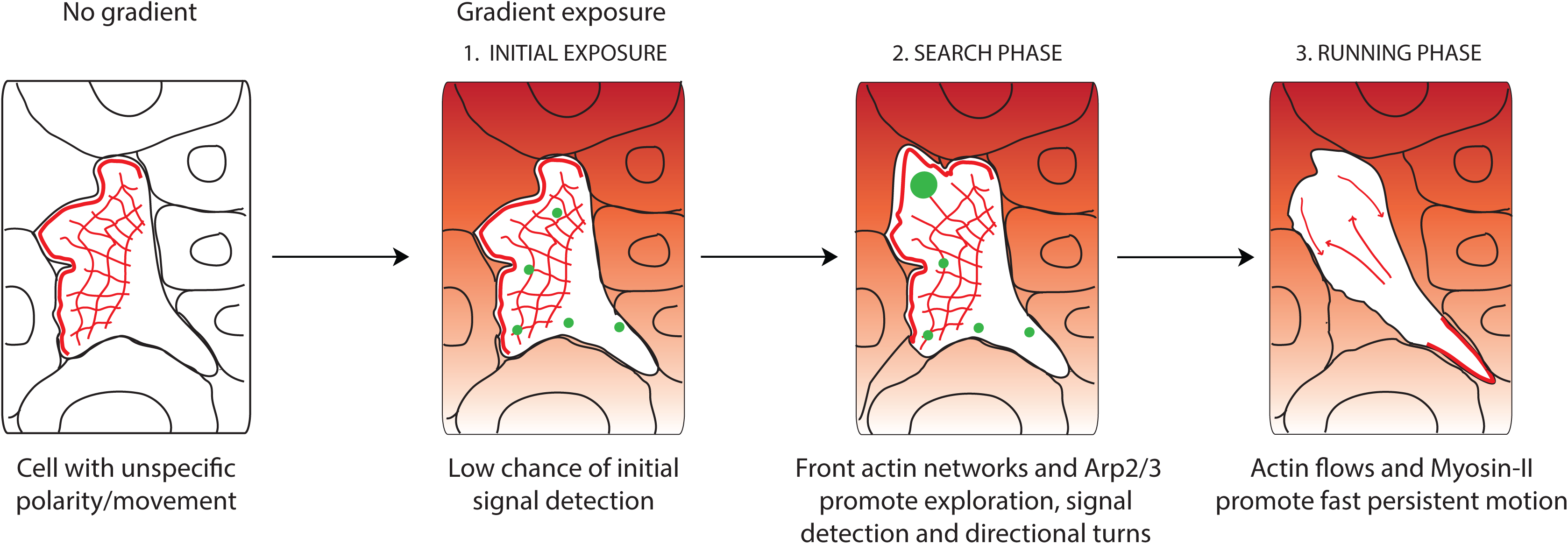
Neutrophils navigate gradients through a search and run strategy. Model for neutrophil gradient sensing *in vivo*. Before encountering the gradient, neutrophils have a pre-established, unspecific polarity and movement. Upon gradient exposure, neutrophils have a low chance of resolving gradient direction without active actin dynamics, therefore a ‘search’ phase is required. During this phase, front actin network expansion via Arp2/3 promotes exploration and signal sampling (illustrated by green spots), culminating in small turns towards the gradient source. Subsequently, cells switch to a ‘running phase’, whereby fast actin flows and Myosin-II contractility enforce fast and persistent motion towards the gradient.

## Discussion

Chemotaxis is fundamental for survival of unicellular and multicellular organisms. Current models are largely based on *in vitro* behaviours of isolated cells or unicellular organisms. How cells navigate in real tissue mazes within a live organism is poorly understood. Here, we used high resolution quantitative imaging and acute chemotaxis assays to derive a more unified model that bridges subcellular dynamics with tissue-scale behaviours. We propose that interstitial navigation requires both leading edge expansion and actin flows for distinct steps in the gradient sensing process. When a wandering cell experiences a new chemical gradient, the first response is to stop and explore (‘searching phase’) rather than steer or accelerate. This is achieved through lateral expansion of front actin networks, which promotes diversification of protrusions and antagonises cell motion. The outcome of these exploratory phases is that protrusions in the direction of the gradient are favoured. Cells then progress to the second stage of the gradient response, whereby actin flows are accelerated promoting fast and persistent motion in the direction of the source (‘running phase’) (Figure 6). We show that protrusive and contractile forces contribute in distinct ways in the gradient response. The search phase is primarily driven by protrusive forces and actin network expansion. On the other hand, the running phase relies on contractile forces that promote actin flows.

Our *in vivo* observations of gradient effects on protrusions are consistent with previous *in vitro* observations. A general premise of current models of chemotaxis is that gradients affect the direction of protrusions, but one matter of debate has been whether these are generated in an oriented fashion in gradients or formed randomly and selectively stabilised (Insall, 2010; Parent et al., 1998; Parent and Devreotes, 1999; Sarris and Sixt, 2015; Servant et al., 2000). Specifically, while strong gradients could induce polarised signalling in immobilised *Dictyostelium amoebae*, shallow gradients rather stimulated random protrusions that were selectively stabilised in the direction of the source based on positive feedback (Insall, 2010; Parent et al., 1998; Parent and Devreotes, 1999). Our data with immobilised neutrophils indicate that gradient exposure leads to a searching pattern of intracellular signalling rather than polarised signalling in the absence of actin dynamics. This suggests that protrusions and actin network expansion are required for establishing polarised signalling rather than represent the outcome of polarised signalling. Thus, gradients encountered by neutrophils *in vivo* are probably too weak to be resolved spatially without active exploration.

Our study nevertheless revealed that leading edge protrusions are only productive if they are succeeded by powerful actin flows that can accelerate and stabilise motion in the direction of the source. In other words, leading edge dynamics only represent the first step in the gradient sensing process. Actin flows are required as a second step, both as a force generating mechanism to accelerate cell motion and as a mechanism of persistence and long-lived memory in cell motion. The role of actin flows as the ultimate output of gradient sensing is consistent with accumulating observations linking actin flows with chemoattractant stimulation and directional persistence (Hons et al., 2018; Maiuri et al., 2015; Yolland et al., 2019). Mechanistically, actin flows can reinforce persistence, through advection of polarity and actin-binding factors (Lange et al., 2016; Maiuri et al., 2015). This provides a cellular memory, which, according to our data, is in the scale of three minutes, given that this is the time frame after gradient exposure that cells persisted in a Myosin-II dependent fashion. This memory mechanism could explain why the first response of cells to a new gradient is to slow down cell movement; to allow the opportunity to reset actin flows (i.e. erase memory from prior movement), renegotiate direction and organise a run in a different direction.

We show differential contributions of diversification of protrusions (via Arp2/3) and contractility (Myosin-II) in the chemotaxis process. Both CK666 and blebbistatin treatments had an effect on speed of locomotion, which was independent of the presence of gradients. In the case of Arp2/3 inhibition, we interpret this as an inability to find paths of least resistance due to compromised diversification of protrusions, as previously shown *in vitro* (Leithner et al., 2016). For Myosin-II inhibition, we reason that impaired contractility directly limits speed due to effects on actin flows and uropod retraction. In regards to gradient sensing, inhibition of Arp2/3 specifically affected the execution of small turns towards the source after the search phase, indicating that protrusion expansion is required for discovering the direction with higher chemical signal. On the other hand, Arp2/3-inhibited cells maintained a bias on speed, despite demonstrating lower global locomotion speed. This indicates that cells can still navigate without diversification of protrusions, albeit not optimally, by biasing speed and persistence towards the source. By contrast, Myosin-II contractility showed a specific defect in persistence and in directional bias on speed, consistent with a requirement for actin flows for the run phase of gradient sensing. We speculate that actin polymerisation through other nucleating factors, such as formins (which extend actin networks in 1D rather than 2D) could suffice to bias actin flows, directional speed and persistence in the forward/backward axis of movement (Vargas et al., 2016; Wilson et al., 2013). It is noteworthy that Arp2/3 inhibition also had an effect on cell persistence in the scale of six but not three minutes. If actin flows provide a cellular memory of a three-minute timescale, this would also be the time scale for each search-and-run cycle. After this time, the cell would ‘forget’ the outcome of its last search and require a reassessment/confirmation of direction through new search. This could explain why diversified protrusions (Arp2/3) influence persistence in gradients in longer timescales.

The gradient sensing model we propose raises several interesting questions. One question is how the transition from searching to running phase is achieved. One possibility is that once cells accumulate signal in one direction (signalling checkpoint model) this stimulates an active switch in contractility and an increase in actin flows. Another possibility is that fast actin flows naturally emerge when actin polymerisation and local actin flows in different parts of the cell become aligned and coordinated in the same direction (Yolland et al., 2019) while they might fail to emerge when initiated in antagonistic directions (emergent behaviour model). We would favour the latter scenario, given that actin flows and cell persistence can also self-organise in unspecific directions in the absence of gradients. Another interesting question is how the two stages of chemotaxis might be affected by tissue geometry. In contrast to a 3D or 2D setting, lateral negotiation of direction and exploratory protrusions are prevented in 1D interstitial channels. This would predict that the relative importance of diversified protrusions in gradient sensing would be lower in such confined environments. It would be interesting to experimentally explore how varied confinement influences the search and run phase of gradient sensing and the resulting motion patterns.

Altogether, our study reveals a unified model of chemotaxis that integrates the role of protrusions and actin flows and explains how cells employ protrusive and contractile forces to interpret gradients within complex tissue settings. Given the fundamental nature of chemotaxis in development, immunity and cancer our findings provide a paradigm of broad physiological relevance.

## Acknowledgements

We thank Hazel Walker and Rob White for comments on the manuscript; Kevin O’Holleran and Martin Lenz of the Cambridge Advanced Imaging Centre, for their support and assistance in this work with two-photon microscopy; Bill Harris, Christine Holt, Ewa Paluch and Kristian Franze groups for spinning disk confocal microscopy equipment; Ronald Germain for intravital microscopy equipment; Tomasz Dyl and Luboš Chvostek for assistance with zebrafish husbandry; Andrei Luchici and Brian Stramer for sharing algorithms for analysis of actomyosin flows; Anna Huttenlocher for the Tg(*mpx*:Lifeact-Ruby) and Tg(*mpx*:PHAKT-EGFP) zebrafish lines; Carl-Philipp Heisenberg for the Tg(*actb1*:myl12.1-eGFP) line; Phil Crosier for the Tg(*lyz*:DsRed2)^nz50^ line. H.P. was supported by a Wellcome Trust PhD grant (105391/Z/14/Z). T.L. was funded by the Max Planck Society and an ERC starting grant (715890). M.S., A.G. and the research were supported by a Medical Research Council Career Development Award (MR/L019523/1), a Wellcome Trust (204845/Z/16/Z), Isaac Newton Trust (12.21 (a)i) and Isaac Newton Trust (19.23 (n)).

## Methods

### General zebrafish procedures

The zebrafish lines used were: Tg(*mpx*:Lifeact-Ruby) (Yoo et al., 2010), Tg(*mpx*:PHAKT-eGFP) (Yoo et al., 2010), Tg(*actb1*:myl12.1-eGFP) (Behrndt et al., 2012) and Tg(*lyz*:DsRed2)^nz50^ (Hall et al., 2007). Zebrafish were maintained in accordance with UK Home Office regulations, UK Animals (Scientific Procedures) Act 1986. Adult zebrafish were maintained under project licenses 70/8255 and P533F2314. Zebrafish were maintained according to ARRIVE guidelines. They were bred and maintained under standard conditions at (28.5 ± 0.5)°C on a 14h-light:10h-dark cycle. Embryos were collected from natural spawning at 4-5 hours post-fertilisation and thereafter kept in a temperature-controlled incubator at 28°C. Embryos were grown in E3 medium, bleached as described in the Zebrafish Book (Westerfield M, 2007) and then kept in E3 medium supplemented with 0.3µg/ml of methylene blue (Sigma-Aldrich, Cat No M9140-25G) and 0.003% 1-phenyl-2-thiourea (Sigma-Aldrich, Cat No P7629-25G) to prevent melanin synthesis. All embryos were used between 2.5-3.5 days post-fertilisation (dpf), thus before the onset of independent feeding. Where indicated, larvae were treated with: 1μM Latrunculin B (Merck, Cat No 428020-1MG), 30nM Leukotriene B4 (Sigma-Aldrich, Cat No L0517-10UG), 40-50μM CK666 (Sigma-Aldrich, Cat No SML0006-5MG), 150μM Blebbistatin (Merck, Cat No 203390-5MG) or DMSO (Sigma-Aldrich, Cat No D8418-100ML).

### Two-photon laser wound and live imaging in zebrafish

For laser wounds, 3dpf larvae were anaesthetised with 0.04% MS-222 (Sigma-Aldrich, Cat No E10521-50G) and mounted onto a glass-bottom plate in 2% low melting agarose (Invitrogen). Agarose-embedded embryos were covered with 2mL E3 medium (supplemented with MS-222). Laser wounding was performed on a two-photon scanning miscroscope (LaVision Biotec TriM Scope II). A tunable ultrafast laser (Insight DeepSee, SpectraPhysics) was tuned to 900nm and the laser power adjusted to approximately 900mW. A square region of interest (ROI) of 20-30µm in width was defined in one focal plane, 10µm below the larva surface, followed by single laser scan across the ROI at a pixel spacing of 240nm and dwell time of 15µs. Confocal stacks were acquired immediately after, using a 25×/1.05 numerical aperture (NA) water-dipping lens. GFP was imaged with 930nm (for PIP3 analysis) and Ruby was imaged with a 1040nm (for Lifeact analysis).

### Mechanical wound and live imaging in zebrafish

For mechanical wounds, 3dpf larvae were anaesthetised with 0.04% MS-222 and their tail-fin was amputated using a sterile surgical scalpel blade (Swann-Morton, 23). Larvae were mounted immediately after wound onto a glass-bottom plate in 2% low melting agarose (Invitrogen). Agarose-embedded embryos were covered with 2mL E3 medium (supplemented with MS-222) and imaged on an inverted PerkinElmer UltraVIEW ERS, Olympus IX81 spinning disk confocal microscope, with a 30×/1.05 NA silicon oil (Olympus) (for PIP3 analysis) or 60×/1.4 NA silicon oil objective (Olympus) (for Myosin-II dynamics), using 488nm laser for GFP excitation and 561nm laser for DsRed excitation. Confocal stacks using a 2µm z-spacing were acquired every 15 seconds (for PIP3 analysis) or every 0.5-2.5 seconds (for Myosin-II analysis).

### General mouse procedures

C57Bl6 mice and C57Bl6-Albino (Tyr^c-2J7c-2J^, JAX000058) mice were purchased from Jackson Laboratories. Mice were maintained in specific-pathogen-free conditions at an Association for Assessment and Accreditation of Laboratory Animal Care-accredited animal facility at the NIAID, National Institutes of Health, and were used under a study protocol from Dr Ron Germain approved by NIAID Animal Care and Use Committee (National Institutes of Health). For intradermal injection experiments, mouse neutrophils were isolated from bone marrow of C57Bl6 mice using a three-layer Percoll gradient of 78%, 69% and 52%. Neutrophils were washed three times with washing buffer (1×Hank’s balanced salt solution, 1% FBS, 2mM EDTA). For fluorescent cell labelling, neutrophils were incubated for 15 minutes with 0.8mM CellTracker Red CMTPX (Invitrogen, Cat No C34552) in 1×HBSS supplemented with 0.0002% (w/v) pluronic F-127 (Thermo Fischer Scientific). Neutrophils were washed four times with washing buffer, before neutrophils (>2×10^6^ cells) were taken up in 1×PBS at a volume of 15-30µL. A volume of 5µL neutrophil suspension was injected intradermally with an insulin syringe (31.5 GA needle, BD Biosciences) into the ventral side of the mouse ear pinnae. In some cases they were co-injected with neutrophils from genetically modified mice that were not analysed here, nor observed to affect behaviour of wild type neutrophils. Recipient mice were always on the C57Bl6-Albino background.

### Two-photon laser wound and live imaging in mice

Two-photon intravital imaging of ear pinnae of anaesthetized mice and laser-induced tissue injury was performed as previously described (Lämmermann et al., 2013). Analysis of mouse neutrophil dynamics was performed on experiments that were referred to, but data not shown, in Lämmermann et al. 2013 (Lämmermann et al., 2013). Two to three hours after injection, mice were anaesthetised and prepared for skin imaging and rested in the heated environmental chamber for 30 minutes before imaging started. Mice were anaesthetised using isoflurane (Baxter; 2% for induction, 1-1.5% for maintenance, vaporised in an 80:20 mixture of oxygen and air) and placed in a lateral recumbent position on a custom imaging platform such that the ventral side of the ear pinna rested on a coverslip. A strip of Durapore tape was placed lightly over the ear pinna and affixed to the imaging platform to immobilise the tissue. Images were captured towards the anterior half of the ear pinna where hair follicles are sparse. Images were acquired using an inverted LSM 510 NLO multiphoton microscope (Carl Zeiss Microimaging) enclosed in a custom-built environmental chamber that was maintained at 32°C using heated air. This system had been custom fitted with three external non-descanned photomultiplier tube detectors in the reflected light path. Images were acquired using a 25×/0.8 NA Plan-Apochromat objective (Carl Zeiss Imaging) with glycerol as immersion medium. Fluorescence excitation was provided by a Chameleon XR Ti:Sapphire laser (Coherent) tuned to 850nm for excitation of CellTracker Red CMTPX dye-labelled neutrophils and second harmonic generation to orient in the connective tissue of the skin. For four-dimensional data sets, three-dimensional stacks were captured every 30 seconds. For focal tissue damage in the ear dermis, the Chameleon XR Ti:Sapphire laser (Coherent) was tuned to 850nm and the laser intensity adjusted to 80mW. At pixel dimensions of 0.14×0.14 µm, a circular region of interest of 15-25µm in diameter (approximately 1-2×10^-6^ mm^3^ in volume) was defined in one focal plane, followed by laser scanning at a pixel dwell time of 0.8µs for 35-50 iterations, depending on the tissue depth of the imaging field of view. Immediately after laser-induced tissue damage, imaging of the neutrophil response was started at typical voxel dimensions of 0.72×0.72×2µm.

### Automated image analysis

#### Extraction of neutrophil trajectories

Analysis of neutrophil trajectories was performed in Imaris v8.2 (Bitplane AG, Zürich, Switzerland) on 2D maximum intensity projections of the 4D time-lapse movies. Analysed trajectories were extracted from the tail fin for mechanical wounds, ventral fin (tracking data in the CHT were excluded) for two-photon ablations for Lifeact analysis or area between eye and ear for two-photon ablations for PIP3 analysis. A track duration threshold of 3 time-points was defined to exclude short-lived tracks. Manual track corrections were also applied where needed. Instantaneous neutrophil coordinates over time (*x,y,z,t*) were exported into Microsoft Excel 2016 spreadsheets files (Microsoft Corporation, Redmond, WA). The latter were imported into MATLAB R2018b (The MathWorks, Inc., Natick, MA) for processing and extraction of results.

#### Definition of mechanical and laser wound perimeter

For laser wounds, the perimeter of the wound was manually defined in MATLAB as a set of points surrounding the largest autofluorescent area around the wound. For mechanical ventral fin wounds, the wound was manually defined in MATLAB as a set of points lining the edge of the amputated tail fin.

#### Neutrophil segmentation

For calculation of mean front and rear actin intensity and Myosin-II flow maps, neutrophil segmentation was achieved with active contours as previously described (Coombs et al., 2019), using custom-written MATLAB scripts. When Tg(*actb1*:myl1.2-eGFP) fish were outcrossed with Tg(*lyz*:DsRed2)^nz50^ fish, segmentation was done first on DsRed neutrophils using active contours, then the generated binary mask was applied on the Myosin-II neutrophils.

#### Definition of actin polarity

For calculation of actin polarity, the separation of a neutrophil into front and rear part was achieved using MATLAB with custom-written scripts. The front and rear parts of neutrophils were defined based on the vector of speed between two successive neutrophil centroids, as calculated from segmentation. A vertical to the speed vector line was defined to separate the neutrophil into two parts. The centre of the line was the centroid of the segmented neutrophil. The front part was formed by grouping pixels with distance from the line higher than 0 while the rear part was formed by grouping pixels with distance from the line below 0. Time plots of actin polarity were performed using the MATLAB function boundedline.m (https://github.com/kakearney/boundedline-pkg).

#### Quantification of PH-AKT dynamics

For calculating PH-AKT mean intensity and standard deviation of PH-AKT mean intensities we used custom-written MATLAB scripts, including MATLAB’s built-in functions *nanmean* and *nanstd*. Contrast was calculated as previously described (Coombs et al., 2019). Cell orientation was calculated as the ratio of mean intensities of ‘towards’ and ‘away’ neutrophil parts. For separation of individual neutrophils into ‘towards’ and ‘away’ segments, a line was defined to connect the geometrical neutrophil centroid with the nearest point of the mechanical tail-fin wound or the centre of the laser wound. A second line, perpendicular to the first line, was defined as passing from the geometrical centroid. The ‘towards’ part was formed by grouping pixels with distance from the line higher than 0 while the ‘away’ part was formed by grouping pixels with distance from the line below 0. Analysis was based on timepoints just before LTB4 addition (-0.25min) or laser wounding (-0.5min) and a time-point within the first 3 minutes after LTB4 addition/laser wound, at which maximal visible change in PH-AKT intensity was observed.

#### Quantification of directional speed, track straightness, angles theta and delta and Myosin-II flows

The calculation of speed versus orientation (directional speed) and of track straightness was achieved using custom-written MATLAB scripts as previously described (Coombs et al., 2019). Neutrophil approach theta and turning delta were calculated using custom-written MATLAB. Instantaneous neutrophil theta was calculated as the angle between the vector of instantaneous speed and the vector that connects the first centroid and the nearest point of the wound perimeter. Instantaneous neutrophil delta was calculated using three time-points and it was defined as the angle between the vector of instantaneous speed between first-second time-frames and the vector of instantaneous speed between first-third time-frames. Scatter plots of angle delta in relation to angle theta were performed using the MATLAB function dcatter.m (https://www.mathworks.com/matlabcentral/fileexchange/8430-flow-cytometry-data-reader-and-visualization). Time plots of speed, theta and delta were performed using the MATLAB function boundedline.m (https://github.com/kakearney/boundedline-pkg).

For calculation of retrograde flow maps, migrating neutrophils were transformed into stationary ones, by subtracting the coordinates of their centroid in each time-frame from the coordinates of their centroid in the first time frame. The calculation of the Myosin-II retrograde flow maps was done using Particle Image Velocimetry (PIV), based on algorithms described previously in (Davis et al., 2015), by choosing the Myosin-II particles with an angle between their speed vector and the vector of neutrophil speed above 145° and below 225° (in the range 0-360°). Correlation coefficients were calculated by custom-written MATLAB scrips, using MATLAB’s built-in cross-covariance function, as previously described (Mueller et al., 2017).

### Statistics

All error bars indicate S.E.M. (Standard Error of the Mean). All *p* values were calculated with two-tailed statistical tests and 95% confidence intervals. Two-tailed paired *t*-test, Wilcoxon matched-pairs signed rank test were performed for pairwise comparisons and ordinary one-way ANOVA with Dunnett’s multiple comparisons test were performed after distribution was tested for normality. Statistical tests were performed in Prism v9.0.1 (GraphPad Software Inc., La Jolla, CA). The statistical test and the *n* number are indicated in the figure legends. The error bars show SEM either across individual embryos (i.e. analysis of population neutrophil persistence-turning) or individual neutrophils pooled from different embryos (i.e. speed versus orientation and orientation versus distance from wound which are step-based analyses, or speed versus time, actin polarity versus time, orientation versus time, turning versus time, correlation coefficient, track straightness, cell contrast, standard deviation, intensity and orientation, which are cell-based analyses).

## Supplementary Figure Legends

**Supplementary Figure 1. Explanatory schemes for acute chemotaxis assays and related analyses.**

(A) Scheme showing two-photon laser wound chemotaxis assay in zebrafish. Left: resting neutrophils at the Caudal Haemopoietic Tissue (CHT). Middle: neutrophils migrating towards the ventral fin (VF) after LTB4 addition. Right: laser wounding in the VF causing redirection of neutrophil movement.

(B) Scheme for angles δ (delta) and θ (theta). Delta is the angle between the speed vector of three successive time-points (t_0_ – t_-1_, t_1_ – t_0_). Theta is the angle between the speed vector (t_1_ – t_0_) and the vector connecting the neutrophil geometrical centre to the closest point of the laser wound perimeter.

(C) Scheme showing two-photon laser wound chemotaxis assay in mice. Neutrophils from bone marrow of C57Bl6 mice were isolated, fluorescently dye-labelled, and then injected into the ear dermis of C57Bl6-Albino mice. Three hours later, intravital imaging started by recording neutrophil dynamics before and after induction of a small tissue lesion in the skin dermis by a two-photon laser beam.

(D) Scheme showing gradient response assay in immobilised neutrophils in zebrafish. Left: migrating neutrophils towards the tail fin (TF). Middle: neutrophils losing polarity and motility after addition of LatB. Right: addition of LTB4 and assessment of signalling response.

**Supplementary Figure 2. Gradients induce a searching pattern of signalling activity in the absence of actin dynamics.**

(A) Scheme of a zebrafish larva indicating the tail-fin (TF) wound (red) and area of imaging (blue). Left: neutrophils in a Tg(*mpx:*PHAKT-EGFP) infiltrating the TF. Red dotted line indicates the TF wound. Middle: neutrophils lose motility post-LatB addition. Right: neutrophils immediately after LTB4 addition. White arrows indicate the neutrophils. Scale bar = 15μm.

(B) Scheme that shows the automated segmentation of neutrophils into two parts, facing towards and away from the wound for analysis of PH-AKT orientation in (C).

(C) Plots of neutrophil PH-AKT contrast, standard deviation, mean intensity and orientation (towards/away), within a time-window of 3 min after LTB4 addition. n=34 cells from 5 larvae. Two-tailed paired *t*-test.

(D) Scheme of larva with indication of laser wounding (LW) (red) and area of imaging (blue). Left: immobilised neutrophils in a Tg(*mpx:*PHAKT-EGFP) larva after LatB addition; Middle: time-point of the LW (red dotted circle). Right: cell response at 1.5 min post-LW. Scale bar = 25μm.

(E) Scheme that shows the separation of cell into two parts, one towards to laser wound and one away from the laser wound. The line that separates the cell is perpendicular to the line that connects the centre of cell and the centre of the laser wound.

(F) Plots of neutrophil PIP3 contrast, standard deviation, mean intensity and orientation (towards/away). n=43 cells from 8 larvae. Two-tailed paired *t*-test.

**Supplementary Figure 3. Effects of Arp2/3 and Myosin-II inhibition on cell speed.** (A,B,C) Neutrophil speed in relation to the cosine theta for DMSO (A), CK666 (B) and blebbistatin (C), for pre- and post-laser wound.(A) DMSO: n=217-1020 cell steps per bin pre-wound, n=179-1734 cell steps per bin post-wound. Data from 9 larvae. (B) CK666: n=131-675 cell steps per bin pre-wound, n=122-825 cell steps per bin post-wound. Data from 5 larvae. (C) Blebbistatin: n=120-427 cell steps per bin pre-wound, n=127-929 cell steps per bin post-wound. Data from 4 larvae.

(D,E,F) Neutrophil speed in relation to time for DMSO (D), CK666 (E) and blebbistatin (F), synchronised with the time of laser wound (LW – 0 min, left) or the time of individual neutrophil beginning of movement (BM – 0 min, right). (D) DMSO: n=17 cells from 8 larvae.

(E) CK666: n=11 cells from 5 larvae. (F) Blebbistatin: n=13 cells from 4 larvae. Mean and SEM are shown.

## Video Legends

**Video 1. Neutrophil migration pre- and post-laser wound in a zebrafish larva. Related to Figure 1 and Figure 2.**

Neutrophils in a Tg(*mpx*:Lifeact-Ruby) zebrafish larva. CHT = Caudal Haemopoietic Tissue. VF = Ventral Fin. LW = Laser Wound. Frame interval is 20 sec and frame rate is 15 fps. Scale bar = 50μm.

**Video 2. Neutrophil migration pre- and post-laser wound in a mouse. Related to Figure 1.**

Migrating neutrophils in the ear dermis of C57Bl6-Albino mouse. Laser wound occurs at 25 minutes, in the skin dermis. Left: Second harmonic generation image (grey) with CTMPX-labelled neutrophils (red). Right: neutrophil channel only. LW = Laser Wound. Frame interval is 30 sec and frame rate is 15 fps. Scale bar = 50μm.

**Video 3. Retrograde Myosin-II flows in zebrafish neutrophils migrating towards a wound. Related to Figure 3.**

(A) Neutrophil in a Tg(*actb1*:myl12.1-eGFP) zebrafish larva, migrating towards the tail-fin wound. Red line denotes the segmented outline of the cell. White arrows indicate the velocity vector for each time-frame. (B) Myosin-II retrograde flows in the reference system of the cell. (C): Myosin-II retrograde flow speed vectors (red). (D): heat-map of flow speeds. Frame interval is 1.93 sec and frame rate is 6 fps. Scale bar = 10μm.

**Video 4. Response of individual neutrophil to laser wound. Related to Figure 4.**

Neutrophil in a Tg(*mpx*:Lifeact-Ruby) zebrafish larva. Laser wound occurs at 6:40 min (not visible in the video, direction is indicated by angle). White arrow indicates the neutrophil to observe. Red arrow indicates Lifeact at the front; green arrow indicates Lifeact at the rear. CHT = Caudal Haemopoietic Tissue. VF = Ventral Fin. LW = Laser Wound. Frame interval is 20 sec and frame rate is 6 fps. Scale bar = 20μm.

**Video 5. PH-AKT distribution in immobilised cells before and after gradient exposure. Related to Supplementary Figure 2.**

(A) Neutrophils in a Tg(*mpx*:PHAKT-EGFP) zebrafish larva initially migrating towards a tail-fin wound (unspecific motion). Time in relation to tail-fin wounding is indicated. Green arrows indicate PIP3 activity post-LTB4. TF Wound = Tail-fin Wound. Frame interval is 15 sec and frame rate is 6 fps. Scale bar = 15μm.

(B) Neutrophils in a Tg(*mpx*:PHAKT-EGFP) zebrafish larva migrating randomly in the head and subsequently exposed to LatB and laser wound. White arrows at 9 min indicate decelerating cells to observe. Arrows post-laser wound indicate cells with PIP3 activity. LW = Laser Wound. Frame interval is 30 sec and frame rate is 6 fps. Scale bar = 50μm.

**Video 6. Effects of inhibitors on actin dynamics and motion patterns. Related to Figure 5 and Supplementary Figure 3.**

Example neutrophils (white arrow) in a Tg(*mpx*:Lifeact-Ruby) zebrafish larva in (A) DMSO, (B) CK666 and (C) blebbistatin, pre- and post-laser wound. CHT = Caudal Haemopoietic Tissue. VF = Ventral Fin. LW = Laser Wound. Frame interval is 20 sec and frame rate is 6 fps. Scale bar = 20μm.

