## Supplementary figures and images for "Interstitial leukocyte navigation through a search and run response to gradients"

### Supplementral Figures

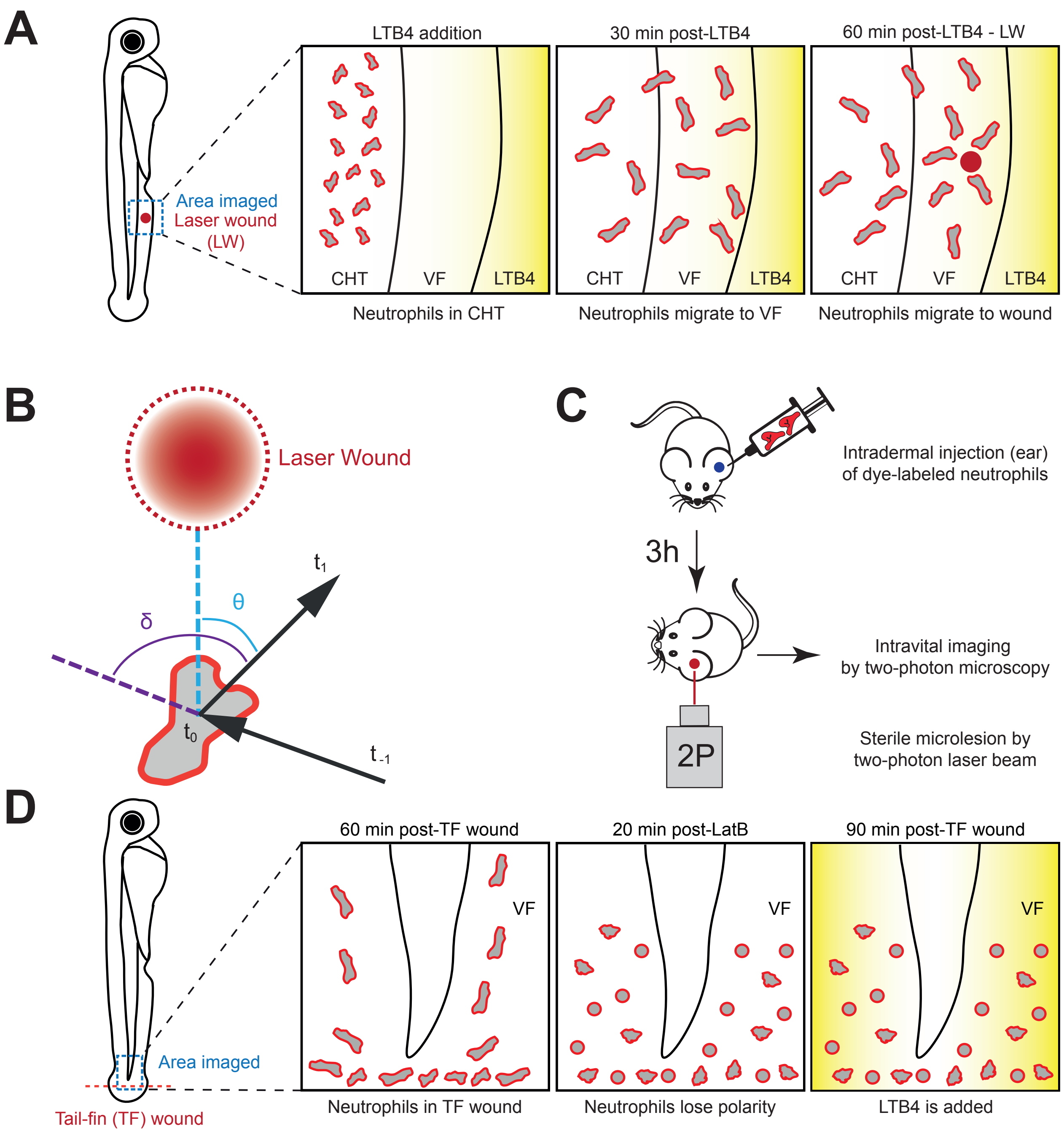

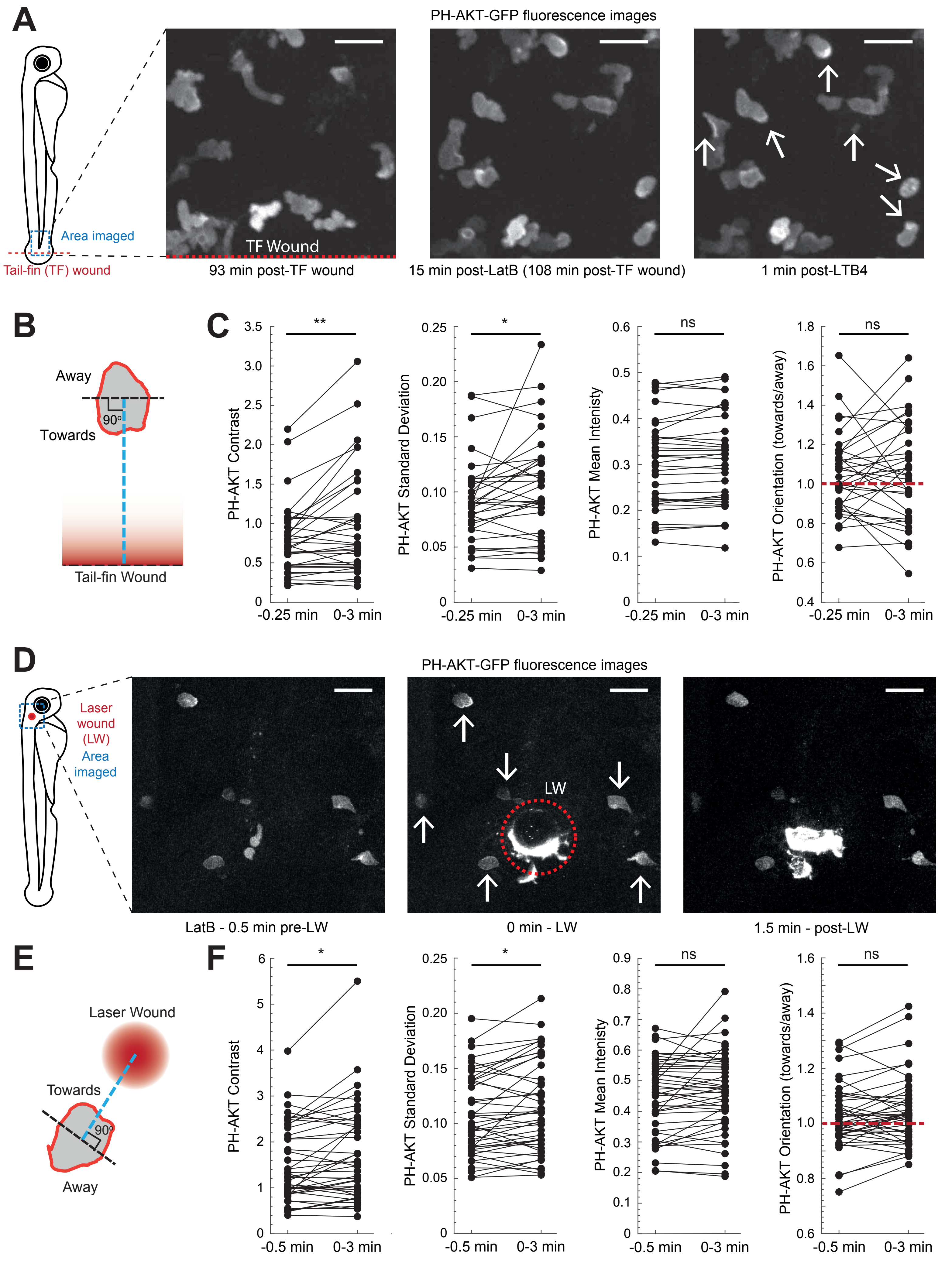

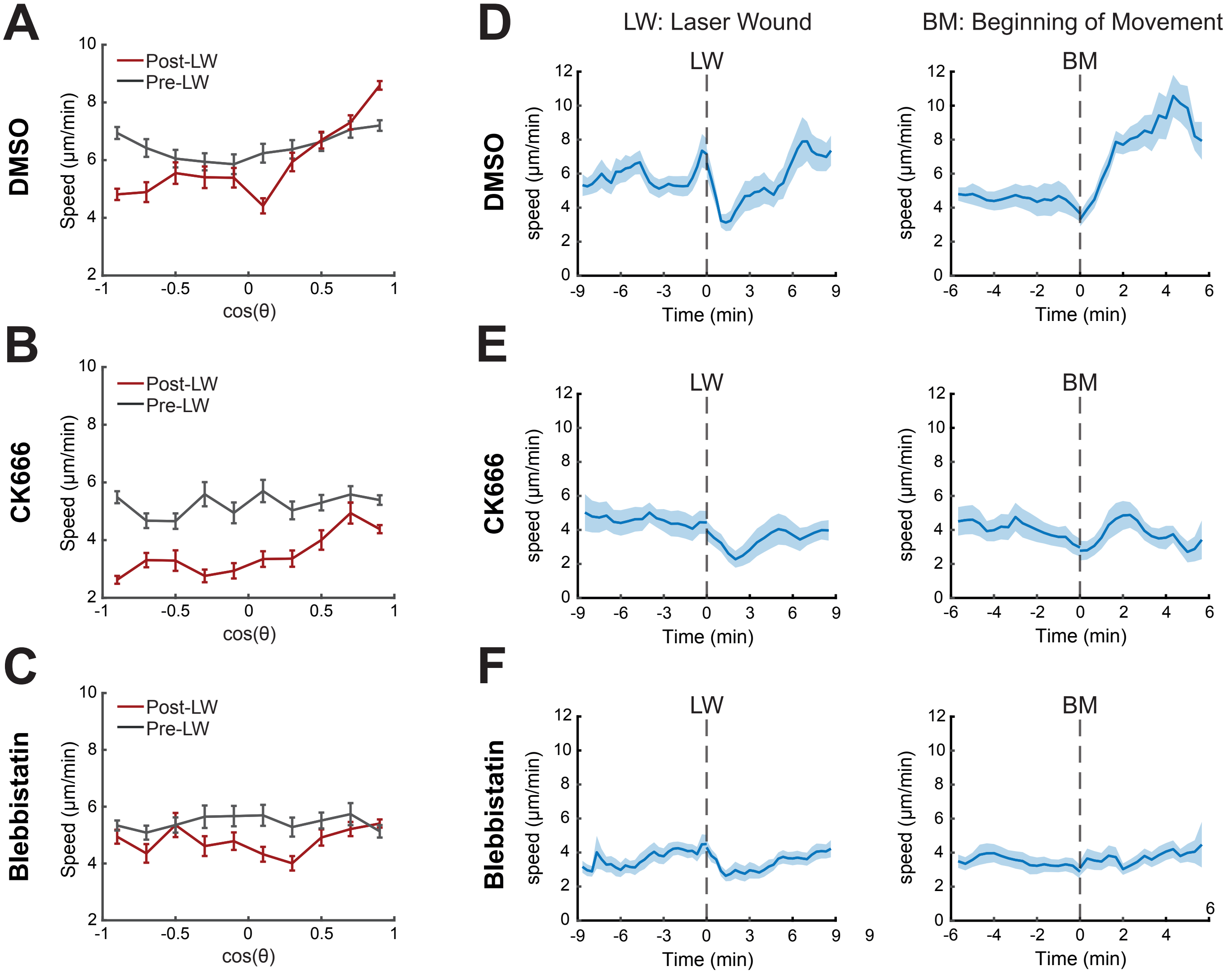
